# Distinct expression patterns of Hedgehog signaling components In mouse gustatory system during postnatal tongue development and adult homeostasis

**DOI:** 10.1101/2023.11.10.566617

**Authors:** Archana Kumari, Nicole E. Franks, Libo Li, Gabrielle Audu, Sarah Liskowicz, John D. Johnson, Charlotte M. Mistretta, Benjamin L. Allen

## Abstract

It is well established that the Hedgehog (HH) pathway regulates embryonic development of anterior tongue taste fungiform papilla (FP) and the posterior circumvallate (CVP) and foliate (FOP) taste papillae, and taste organ maintenance and regeneration in adults. However, there are knowledge gaps in determining HH signaling regulation in postnatal taste organ differentiation and maturation. Importantly, the HH transcription factors GLI1, GLI2 and GLI3 have not been investigated in early postnatal stages; and, the receptors PTCH1, GAS1, CDON and HHIP, required to either drive HH pathway activation or antagonism, remain unexplored. Using *lacZ* reporter mouse models, we mapped expression of the HH ligand SHH, receptors, and transcription factors in FP, CVP and FOP in early and late postnatal and adult stages. In adults we also studied the soft palate, and the geniculate and trigeminal ganglia which extend afferent fibers to the anterior tongue. *Shh* and *Gas1* are the only components that were consistently expressed within taste buds of all three papillae and the soft palate. In the first postnatal week, we observed a broad expression of HH signaling components in FP and adjacent, non-taste filiform (FILIF) papillae in epithelium or stroma and tongue muscles. Remarkably, we observed elimination of *Gli1* in FILIF and *Gas1* in muscles, and downregulation of *Ptch1* in lingual epithelium and of *Cdon*, *Gas1* and *Hhip* in stroma from late postnatal stage. Further, HH receptor expression patterns in CVP and FOP epithelium differed from anterior FP. Among all the components, only known positive regulators of HH signaling, SHH, *Ptch1*, *Gli1* and *Gli2*, were expressed in the ganglia. Our studies emphasize differential regulation of HH signaling in distinct postnatal developmental periods and in anterior versus posterior taste organs, and lay the foundation for functional studies to understand the roles of numerous HH signaling components in postnatal tongue development.

## Introduction

The tongue houses taste buds (TB) in three different taste papillae at three different locations: fungiform papilla (FP) on the anterior two thirds, circumvallate papilla (CVP) on the posterior third mid-dorsum and foliate papilla (FOP) bilaterally on the posterior sides. These papilla types and resident taste buds develop with different time courses in mammalian species (1–3). In sheep for example, extensively studied, the TB make a fetal appearance, continue development *in utero* and have extended postnatal maturation (4). In rat and mouse, the lingual taste papillae emerge in the embryo but although initial TB-like cell clusters are seen before birth, TB *per se* do not emerge until postnatal stages (5–7). However, both papillae and TB continue to develop during early postnatal stages and reach maturity in the anterior and posterior regions by the third and sixth postnatal weeks (1, 8–10). A mature FP harbors a single TB, while a CVP and multiple rows of FOP house numerous TB tightly associated with each other (**Fig 1A**). Anterior tongue and FP receive innervation from chorda tympani and lingual nerve fibers and posterior tongue and CVP are innervated by glossopharyngeal nerve (**Fig 1A)**. Interestingly, FOP anterior ridges receive chorda tympani nerve fibers and posterior ridges are innervated by the glossopharyngeal nerve (**Fig 1A)** (11–13). The distinct location, morphology, and innervation of three lingual taste papillae may lead to their different trajectories of embryonic and postnatal development.

**Fig 1.**
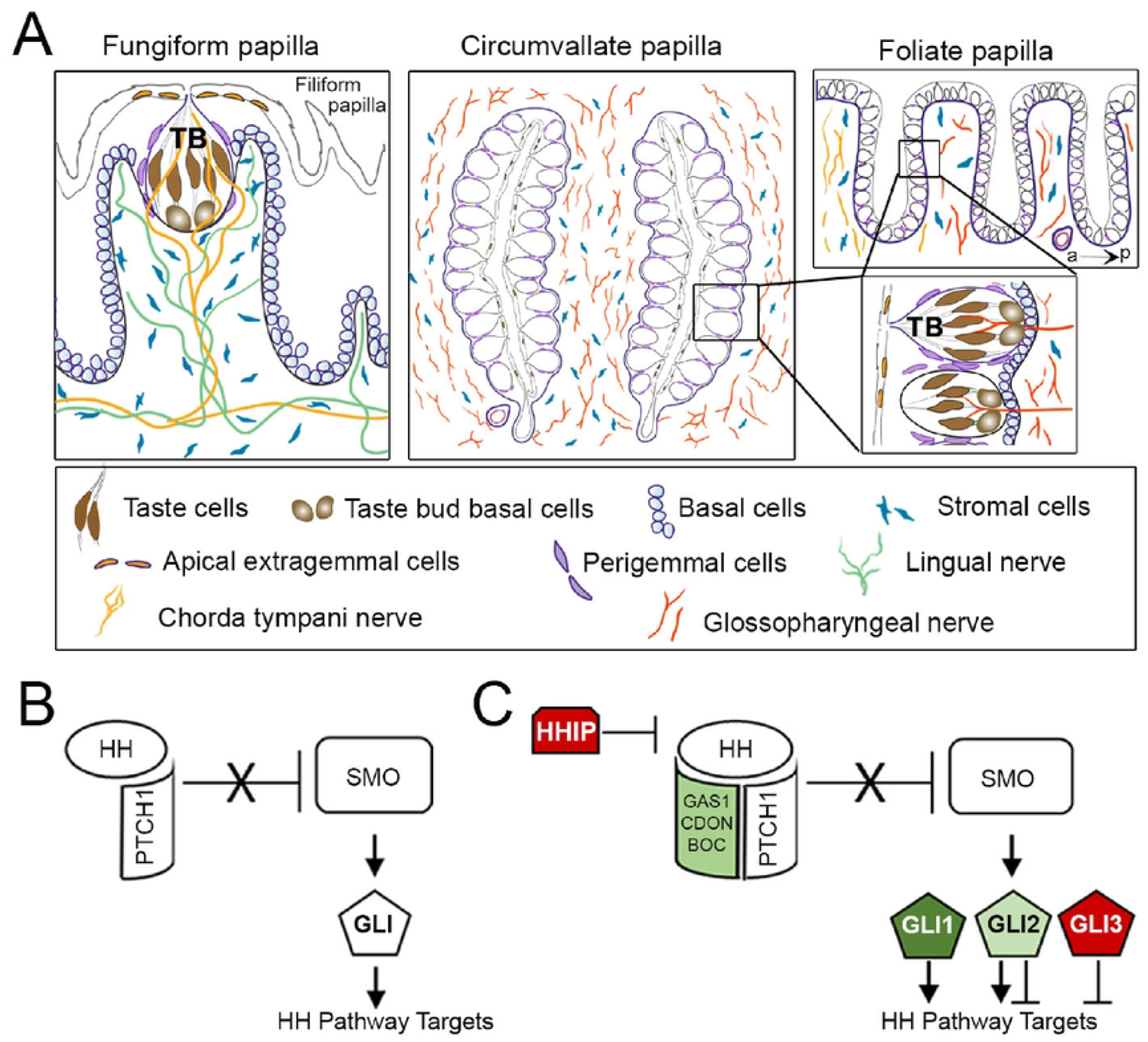
Schematic of lingual taste papillae and Hedgehog (HH) signaling. (A) Illustrations of tissue architecture for the fungiform (FP), circumvallate (CVP) and foliate (FOP) papillae. FP is presented in a sagittal section, surrounded by spinous, non-taste filiform papilla. The CVP and FOP are illustrated in a horizontal section (to the dorsal tongue surface), and boxes refer to magnified taste bud (TB) distribution along the epithelium. FP harbors a single TB, while CVP and FOP can hold numerous TB adjacent to each other. Each TB is composed of taste cells and TB basal cells, and is surrounded by perigemmal cells. All the papillae comprise basal cells in the epithelium, extragemmal cells on the apical surface and stromal cells in the connective tissue core. Gustatory nerves, chorda tympani and glossopharyngeal innervate FP and CVP, respectively. FOP has parallel rows of multiple ridges and the anterior (a) ridges receive chorda tympani while the posterior (p) receives glossopharyngeal nerve fibers. (B) HH ligand binds to the receptor, PTCH1, resulting in de-repression of SMO and the initiation of a downstream signaling cascade that culminates in the modulation of GLI transcription factor activity and HH target gene expression. (C) HH signaling is regulated at the cell surface by the co-receptors, GAS1, CDON and BOC (green, activator), and the secreted antagonist HHIP (red, repressor). Transcriptional readouts of HH signaling are controlled by three GLI proteins: GLI1 (green, activator), GLI2 (light green, activator>repressor), and GLI3 (red, repressor>activator).

In our current study, we use mouse models. The number of mouse FP taste cells increases rapidly by postnatal day 7 (P7), more slowly between P7 and P14, and finally reaches a steady level by P21 (10, 14, 15). In contrast, the number of rodent CVP and FOP TB increases threefold between P7 and P45 and reaches a maximum at P90 (1, 10, 16). In addition, formation of taste pores, remodeling of gustatory innervation and enlargement of tongue muscles also occurs during the initial postnatal weeks (10, 16–18). These various and extreme changes must be tightly coordinated to create a stereotypical, functionally mature taste organ.

Hedgehog (HH) signaling is an essential regulator of embryonic patterning (19–21), postnatal organ development (22, 23) and homeostasis (24) of several organs. We have previously shown the role of epithelial HH/GLI signaling in the maintenance of adult taste organs (25), and that HH signaling through Smoothened (SMO) also regulates papilla/TB homeostasis and neural responses (26–28). HH signaling is also active and vital during tongue development (5, 7, 29, 30). However, few studies have investigated HH signaling regulation in the anterior tongue during the initial postnatal weeks when the dynamic papillae and TB are continuously growing (8, 10, 15). Further, HH signaling regulation in posterior CVP and FOP remain to be investigated. Thus, identification of signaling component activities during postnatal lingual development and maturation is essential to understand HH regulatory roles.

In the basic pathway it is known that HH signaling initiates through ligand binding to the canonical receptor Patched 1 (PTCH1), which relieves inhibition of SMO, which can then mediate downstream signaling with the modulation of GLI transcription factor activity for transcription of HH pathway targets (**Fig 1B**) (31–33). Notably, HH pathway activation also requires ligand interactions with the co-receptors CDON, BOC and GAS1 (34), whereas the pathway is inhibited by the HH antagonist HHIP interaction with the ligand (**Fig 1C**) (35, 36). Furthermore, HH transcription requires the combined use of three different GLI proteins. GLI1 is a transcriptional target and encodes an exclusive activator of the HH pathway; GLI2 is the major activator; and GLI3 acts principally as a repressor in HH signal transduction (**Fig 1C**) (37). The field of taste research has mainly focused on the ligand, sonic HH (SHH), and the target gene, *Gli1*, while PTCH1, GLI2 and GLI3 have received less attention. Further, the obligatory membrane receptors GAS1, BOC and CDON and the HH antagonist HHIP remain largely unexplored. Our recent studies with HHIP (27) emphasize that these less studied HH pathway components might not be ‘secondary’ signaling elements but rather key signaling molecules that regulate organ integrity and thus need addressing.

As FP, CVP and FOP are distinct in their development and structural organization, their signaling regulation might be different. Thus, we investigated the expression pattern of HH signaling components in anterior FP and posterior CVP and FOP tissue regions: TB, basal, perigemmal, apical extragemmal and stromal cells (**Fig 1A**). We studied the ligand *Shh*, membrane receptors *Ptch1, Gas1, Cdon* and HH antagonist *Hhip,* and transcriptional factors *Gli1, Gli2* and *Gli3* in developing FP, CVP and FOP at the early postnatal stages between P3 and P10 (termed P7), and compared the pattern with that of mature taste papillae in adult mice aged eight weeks or older. We also analyzed anterior FP during a later development stage between P21 and P31 (termed P28). In addition, we mapped HH pathway components in the adult soft palate, and geniculate and trigeminal ganglia with soma for chorda tympani and lingual nerve fibers, respectively, in the anterior tongue. We have used X-gal staining of *lacZ*-reporter mouse models or immunostaining to map the pathway components. The data reveal unique expression of the positive regulators of HH signaling activity, *Ptch1, Gas1, Cdon, Gli1* and *Gli2* in FP, CVP and FOP, and soft palate. However, differences are noted in the expression pattern of several HH signaling components in the anterior tongue, which is broader in the early postnatal stages as compared to adulthood. Further, SHH, receptor *Ptch1* and transcription factors, *Gli1* and *Gli2* are the only components of the HH pathway observed in both geniculate and trigeminal ganglia. Overall, our data suggest a shift in HH pathway regulation of taste organ after conclusion of postnatal maturation of tongue.

## Materials and Methods

### Mice

All animal use and care procedures were performed according to the guidelines of the National Institutes of Health and approved protocols of the University of Michigan and Rowan University Institutional Animal Care and Use Committee. Male or female mice, from the first postnatal week through adult stages were used.

### Mouse models/strains

#### *lacZ* reporters

Mice carrying *lacZ* alleles for the ligand (*Shh^lacZ/+^*, MGI:2678342) (38, 39), HH-receptor *Ptch1* (*Ptch1^lacZ^*^/+^, Jackson Laboratories strain:003081) (40), HH co-receptor *Cdon* (*Cdon^lacZ/+^*) (41), HH antagonist *Hhip* (*Hhip^lacZ/+^*, Jackson Laboratories strain:006241) (42), HH target gene/responding *Gli1* (*Gli1^lacZ^*^/+^) (Jackson Laboratories strain:008211), HH transcriptional activator *Gli2* (*Gli2^lacZ/+^*, Jackson Laboratories strain:007922) (43) and transcriptional repressor *Gli3* (*Gli3^lacZ/+^*) (44) were maintained on a mixed 129S4/SvJaeJ/C57BL6/J backgrounds. HH co-receptor *Gas1* (*Gas1^lacZ/+^*) (20) was maintained on a C57BL6/J background. For each reporter, observations were made in at least 3 mice.

#### RFP reporter

Mice carrying tamoxifen inducible expression of RFP in *Shh*-expressing cells and their progeny (*Shh^CreERT2^;R26^RFP^*). *Shh^CreERT2^* (Jackson Laboratories strain:005623); *R26^RFP^* (Jackson Laboratories strain:007914) double transgenic animals were given 400 mg of tamoxifen/kg in Teklad Global Diet (Harlan) daily for 30 days.

### Tissue dissection and processing

Tongues on mandibles were collected between postnatal day (P) 3 and P10 (termed P7), between P21 and P31 (termed P28) and adult (more than 8 weeks of age) stages. Soft palate and the ganglia (geniculate, GG and trigeminal, TG) were dissected at adult stages. All the tissues were fixed for 2h at 4°C in 4% paraformaldehyde in PBS. Fixed tongues were cut to obtain anterior two thirds to include fungiform (FP) and filiform (FILIF) papillae and the posterior third to include circumvallate (CV) and foliate (FOP) papillae. After fixation, all tissues were cryoprotected overnight with 30% sucrose in PBS and embedded in O.C.T. compound (Tissue-Tek, Sakura Finetek) for X-Gal staining or immunostaining as described previously (26, 27). Serial sagittal sections (FP), horizontal sections (CV, FOP, GG and TG) or coronal sections (soft palate) were cut at 10 μm for immunostaining (26, 27).

### X-Gal staining

Tissue sections were incubated in X-Gal solution for 4h-48h. X-gal staining was followed by immunostaining of TB cells (K8) or epithelium (Ecad) in FP sections and ligand (SHH) in GG and TG sections.

### Immunostaining

Immunoreactions were performed as described previously (26, 27). Briefly, tongue or ganglion sections were air dried, rehydrated, blocked (in 10% normal donkey serum, 0.3% Triton-X in PBS-X), and incubated overnight at 4°C with primary antibodies. On the next day, slides were washed and incubated with appropriate secondary antibodies for 1–2 h at room temperature in the dark. Primary antibodies were goat anti-SHH (AF464, 0.1 μg/mL; R&D Systems); rat anti-keratin 8 (TROMA-1, 1:1,000; Developmental Studies Hybridoma Bank); goat anti–E-cadherin (AF748, 1:5,000; R&D Systems) and rabbit anti-RFP (600-401-379, 1:1,000; Rockland). For SHH in tongue sections, the heat-induced antigen-retrieval method was used.

### Imaging

Tissue section images were acquired with a Nikon Eclipse 80i microscope and Nikon DS Ri2 camera system or an automated Keyence BZ-X810 microscope. Photomicrographs were adjusted for brightness and contrast in parallel across one Fig and assembled with Adobe Photoshop.

## Results

### Expression pattern of ligand *Shh* and membrane receptors *Ptch1*, *Gas1*, *Cdon* and *Hhip* in the anterior tongue

In **Fig 1A** specific cell types in FP are illustrated. Our focus here is to compare expression patterns of HH pathway components across three developmental stages (early postnatal at P7, later at P28 and mature beyond 8 weeks of age). *Shh* ligand is present in FP TB, as observed with X-gal staining of the *Shh^lacZ^* reporter mouse model at P7, P28 and adult tongues (**Fig 2A-C**). We do not observe ligand expression in nerves with X-gal staining as reported with transgenic reporter models for *Shh* (26, 45–47). The data show stable *Shh* expression within TB, including TB basal cells, from early postnatal through adult tongues and align with previous studies using the SHH antibody detection method (25–27).

**Fig 2.**
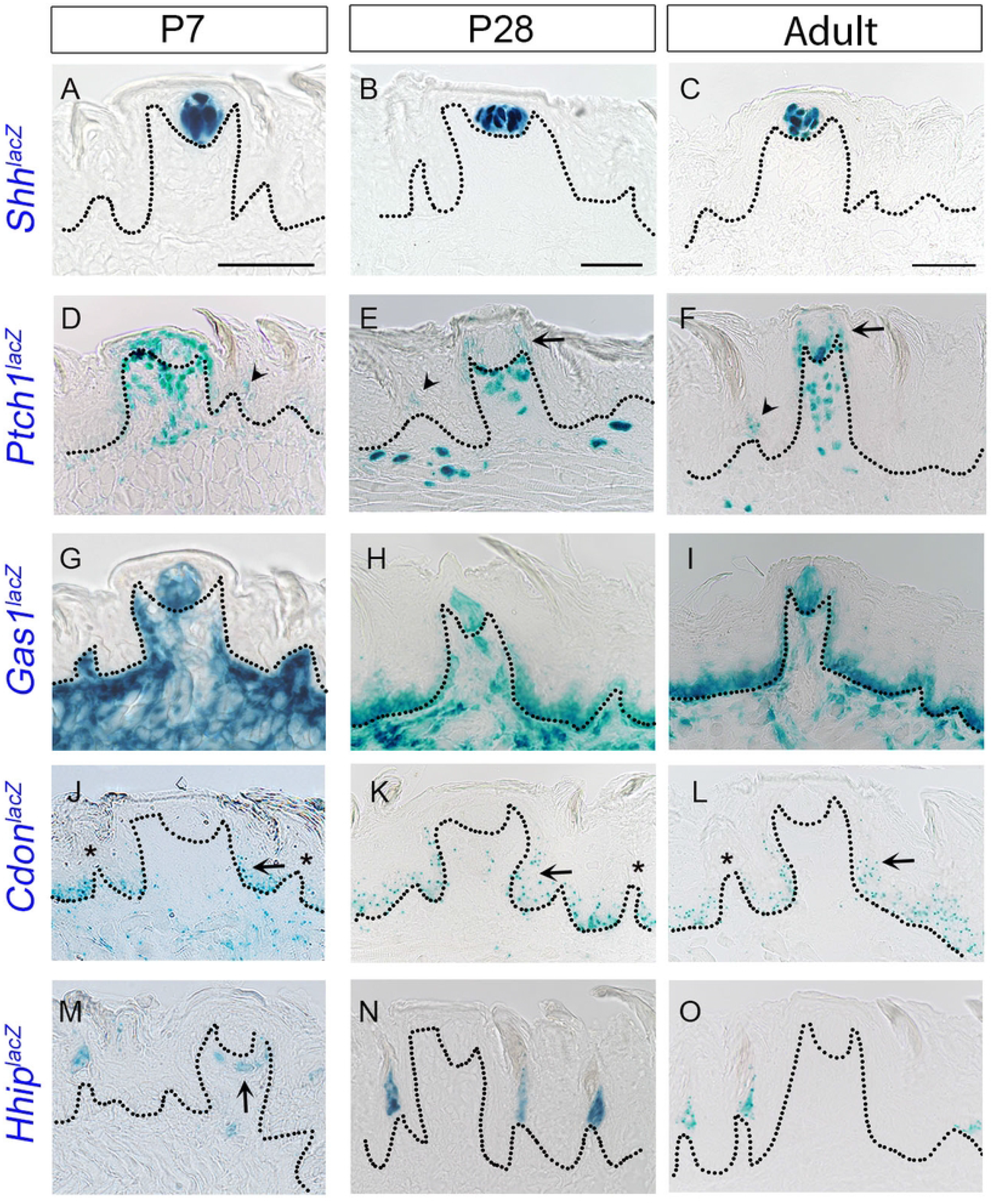
Novel expression of HH pathway receptors in the anterior tongue from early postnatal to adult stages. (A-O) X-gal staining using *Shh^lacZ^*^/+^, *Ptch1^lacZ/+^*, *Gas1^lacZ/+^*, *Cdon^lacZ/+^* and *Hhip^lacZ/+^* reporter mice at the P7, P28 and adult stages. *Shh* ligand is expressed within taste bud (TB) at P7 (A), P28 (B) and adult tongue (C). *Ptch1^lacZ^* is expressed in the FP epithelium at all stages (D-F), but is predominant in the apical region (E,F arrows) from late postnatal to adulthood. In addition, *Ptch1^lacZ^* is expressed in filiform papillae (FILIF) epithelium (D-F, arrowheads) and tongue stromal cells. *Gas1^lacZ^* is expressed in TB at all stages (G-I), extensively in stroma at P7 (G) and induced expression in epithelium from P28 while having less extensive stromal expression (H,I). *Cdon^lacZ^* is observed in the entire lingual epithelium, within FP, it is concentrated in the lower half of the epithelium (J-L, arrows) and within FILIF, it does not overlap with *Ptch1* locations (J-L, asterisks). Stromal *Cdon^lacZ^* expression is also decreased from P7 to adult stages (J-L). The HH antagonist *Hhip^lacZ^* is observed in FILIF at all the stages (M-O). Remarkably, *Hhip^lacZ^* expression is also observed in the FP connective tissue core at P7 (M, arrow). Dotted lines outline the epithelium in all images. Scale bars (50µm) in A,B and C apply to respective column images.

The HH membrane receptor *Ptch1* is expressed in both taste FP and filiform papillae (FILIF) from P7 to 8 weeks of age, as indicated by X-gal staining (**Fig 2D-F**). In FP at P7, *Ptch1^lacZ^* expression is observed in TB, perigemmal, apical extragemmal, basal and stromal cells (**Fig 2D**). At P28, there is no *Ptch1^lacZ^* expression within TB and expression in basal epithelial cells is limited to apical regions of FP, next to perigemmal cells (**Fig 2E**, arrow), while perigemmal, apical extragemmal and stromal cell expression are maintained (**Fig 2E**). This restricted basal cell pattern is also observed in the adult stage (**Fig 2F**, arrow). Upon investigating an earlier stage, P19, we again find a reduction in *Ptch1^lacZ^* expression in FP basal cells and loss of expression within TB (**Fig S1A**). The data clearly indicate a shift in *Ptch1* expression in FP epithelium between P7 and P19. *Ptch1^lacZ^* expression in adult FILIF was reported in the anterior epithelial face in cells of the basal and suprabasal layers (27). We observed that the pattern is maintained from early postnatal stages to adulthood (**Figs 2D-F**, arrowheads; **S1B**).

X-gal staining of *Gas1^lacZ^* mouse tongue revealed expression in FP TB from P7 to adult stages (**Fig 2G-I**). Importantly, at P7, there is no epithelial *Gas1^lacZ^* expression (**Figs 2G**;**S1C**, Ecad). However, *Gas1^lacZ^* expression in lingual epithelium is observed on and after P21 (**Figs 2H,I; S1D,E**). In FP, *Gas1^lacZ^* expression is virtually absent in perigemmal and apical extragemmal cells, and the pattern is maintained through the adult stage (**Fig 2G-I**). Further, we find extensive *Gas1^lacZ^*+ cells in the lingual stroma (**Fig 2G**) and muscles (**Fig S1C**) at or before P7. However, stromal expression declines while expression in muscle ceases on and after P21 (**Figs 2H,I**; **S1D,E**). We find that the expression pattern of HH receptor *Gas1^lacZ^* changed between P7 and P21, as indicated by a reduction in stromal cells, elimination in muscle cells and simultaneous emerging expression in lingual epithelial cells.

Another membrane HH co-receptor, *Cdon^lacZ^*, is expressed in the entire lingual basal epithelium (but not in the apical half of FP wall) from P7 to the adult stage (**Fig 2J-L).** This is unlike other HH receptors, *Ptch1* and *Gas1,* that change their lingual epithelial expression from early postnatal stages to adulthood. Intriguingly, in FP, *Cdon^lacZ^* expression is predominant in the basal lower half of FP (**Fig 2J-L**, arrows) and in FILIF does not overlap with *Ptch1* expression (**Fig 2J-L**, asterisks). Although epithelial *Cdon^lacZ^* expression in FP and FILIF is retained, its expression in overall tongue stromal cells showed reduction at eight weeks of age as compared to P7 or P28 (**Figs 2J-L; S1F,G**). There is no *Cdon ^lacZ^* expression in TB, perigemmal and apical extragemmal cells in any of the tongue developmental stages (**Figs 2J-L; S1F,G**).

Similar to our recent study (27), HH antagonist *Hhip^lacZ^* expression is observed in FILIF apical cells, consistently from P7 to adult stages (**Fig 2M-O**). We do not observe *Hhip^lacZ^* expression in TB, perigemmal and apical extragemmal cells (**Fig 2M-O**). Remarkably, we observe expression of *Hhip^lacZ^* in a few stromal cells within FP connective tissue core concentrating below the TB and in tongue stromal cells at P7. Stromal *Hhip* expression is noted at P12 (**Fig S1H** arrows) but not at P28 and adult stage (**Figs 2N,O; S1I**).

Overall, data indicate that all membranous HH-receptors are expressed in the anterior tongue with partial to no co-expression suggesting a distinct and non-overlapping function in the development and maintenance of taste and non-taste organs. Intriguingly, we observed shifts in the unique expression patterns of *Ptch1*, *Gas1* and *Hhip* in the early postnatal period primarily between P7-P21 suggesting a shift in function or a multi-functional role. Among the four HH membrane receptors studied here, only *Gas1^lacZ^* is consistently present in the TB.

### Transcription factor *Gli1* expression changes during initial postnatal weeks, whereas *Gli2* and *Gli3* expressions remain stable in anterior tongue

As reported earlier (9, 25–27, 48), *Gli1^lacZ^* expression is seen in FP basal, perigemmal, apical extragemmal and stromal cells and is maintained throughout postnatal and adult stages (**Fig 3A-C**). However, we additionally observed *Gli1^lacZ^+* cells in FILIF at P7 (**Fig 3A**, arrows), which are not seen after P21 (**Fig 3B,C**). To determine when *Gli1* expression becomes restricted in the FP, we stained a later developmental timepoint (P14) in *Gli1^lacZ^* mouse tongues. We found that 50% of FP lost *lacZ* expression in neighboring FILIF (**Fig S1J**). After P21, *Gli1^lacZ^*+ cells are observed only in FP (**Fig 3B,C)**. Further, the expression is noted in basal and a few suprabasal cells at P7 (**Fig 3A**, arrowheads) but is restricted to FP basal cells only after P21 (**Fig 3B,C**).

**Fig 3.**
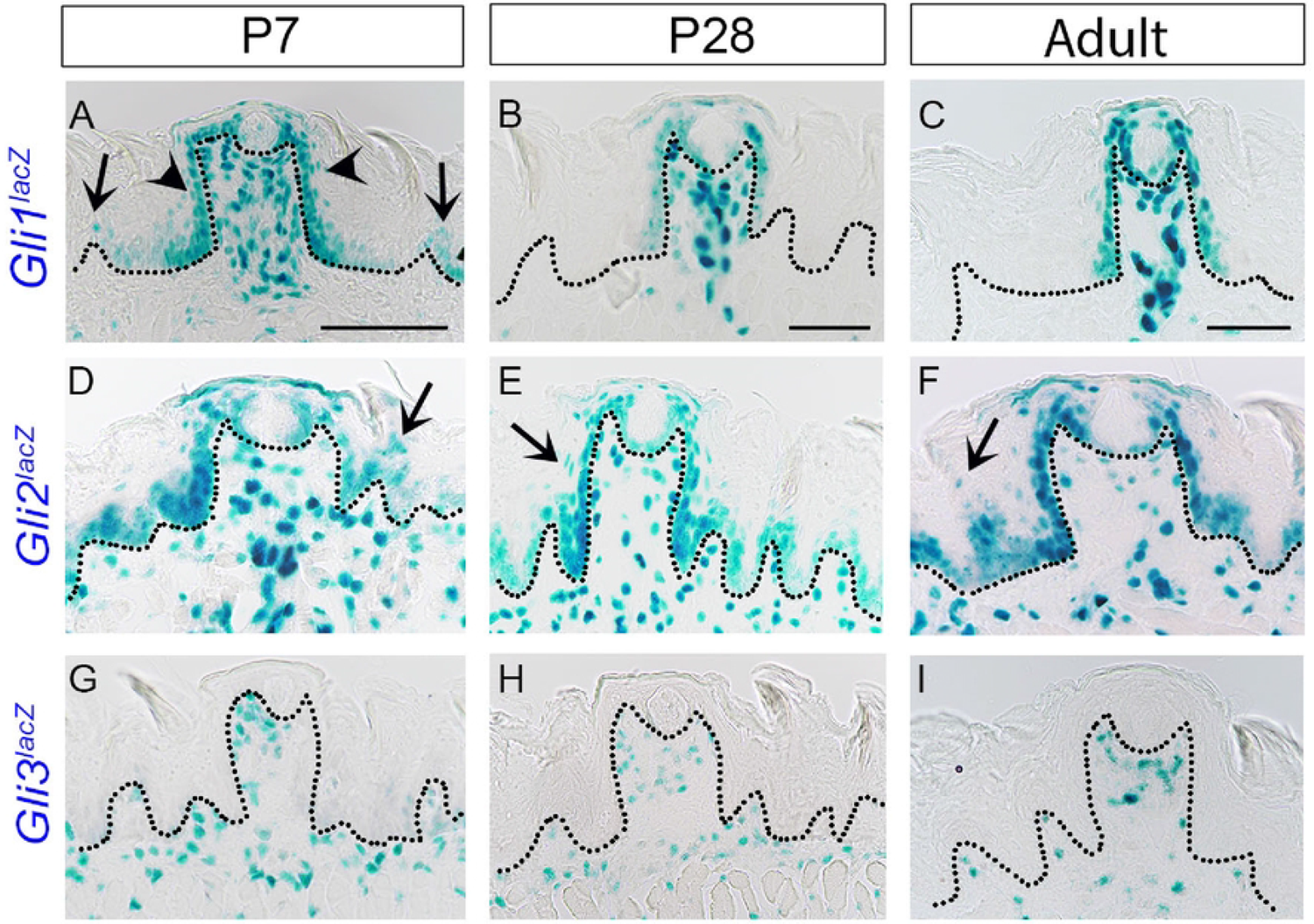
HH pathway transcription factors expression in the anterior tongue from early postnatal to adult stages. (A-I) X-gal staining of *Gli1^lacZ/+^*, *Gli2^lacZ/+^*and *Gli3^lacZ/+^* reporter mice at the P7, P28 and adult stages. *Gli1^lacZ^* is expressed in FP basal, perigemmal, apical extragemmal and stromal cells (A-C). In addition, at P7, *Gli1^lacZ^* is observed in filiform papillae (A, arrows) and FP suprabasal epithelial layers (A, arrowheads). *Gli2^lacZ^* expression is observed in entire lingual basal epithelium, suprabasal layers (D-F, arrows), perigemmal, apical extragemmal and stromal cells in (D-F). In contrast to *Gli1^lacZ^* and *Gli2^lacZ^*, *Gli3^lacZ^* is only present in lingual stromal cells (G-I). Dotted lines outline the epithelium in all images. Scale bars (50µm) in A,B and C apply to respective column images.

The transcription factor, *Gli2* expression is observed in the entire lingual basal, perigemmal, apical extragemmal cells, including both FP and FILIF; and also in stromal cells from P7 to adult stages in *Gli2^lacZ^* mice (**Fig 3D-F**). In addition, a few suprabasal epithelial cells of both FP and FILIF show *Gli2^lacZ^* expression (**Fig 3D-F**, arrows). Stromal expression of *Gli2^lacZ^* cells in the tongue is extensive as compared to *Gli1^lacZ^* cells throughout all stages (**Fig 3D-F**). There is no overt change or shift in the expression pattern of *Gli2^lacZ^* in the adult tongue as compared to the early postnatal stages.

X-gal staining of the *Gli3^lacZ^* mouse tongue revealed positive cells throughout the lingual stroma in both FP and FILIF, which remained unaltered from P7 to adult stages (**Fig 3G-I**). Unlike other *Gli* transcription factors, *Gli3* is not expressed in any of the epithelial (basal, perigemmal and apical extragemmal) cells. We found faint *Gli3^lacZ^* expression in a few TB in the adult stage after 48 hours of X-gal staining). A previous study in adult mice reported *Gli3*+ TB cells by *in situ* hybridization using digoxigenin-labeled *Gli3* RNA probes (49).

The data suggest somewhat overlapping patterns of epithelial expression of *Gli1* and *Gli2* in FP and FILIF during initial postnatal stages. Similar *Gli1* and *Gli2* expression in FP are retained in late postnatal through adult stages. In contrast, *Gli3* is observed only in stromal cells. Whether there is any overlap between *Gli* stromal expressions was not studied.

### Mapping ligand *Shh* and membrane receptors *Ptch1, Gas1, Cdon* and *Hhip* expression in posterior circumvallate and foliate papillae at early postnatal and adult stages

Illustrations for CVP and FOP structure and cell types in the horizontal plane are shown in **Fig 1A**. Based on the developmental trajectory of posterior CVP and FOP, we studied the initial postnatal week and mature adult stages for HH pathway component expression (**Figs 4,5)**. Immunostaining with SHH antibody or X-gal staining of the *Shh^lacZ^* reporter mouse model at P7 showed the ligand expression in CVP and FOP TB (**Figs 4A,C**, K8+; **S1K**). To visualize *Shh* expression in nerves, we used tissues from *Shh^CreER^;RFP* to determine whether nerves are labeled in adult CVP and FOP. After 30 days of tamoxifen treatment in *Shh^CreER^;RFP*, we analyzed CVP and FOP sections and observed *Shh* expression in nerve fibers surrounding the CVP walls and all FOP ridges in the connective tissue core (**Fig 4B,D**, arrows) in addition to TB. Another study using *Shh;mTmG* model also showed *Shh*+ fibers within CVP TB (47). CVP and the posterior-most folds of the FOP are innervated by glossopharyngeal nerves (**Fig 1A**), that have soma in the petrosal ganglion. It is not known whether the petrosal ganglion expresses *Shh*.

**Fig 4.**
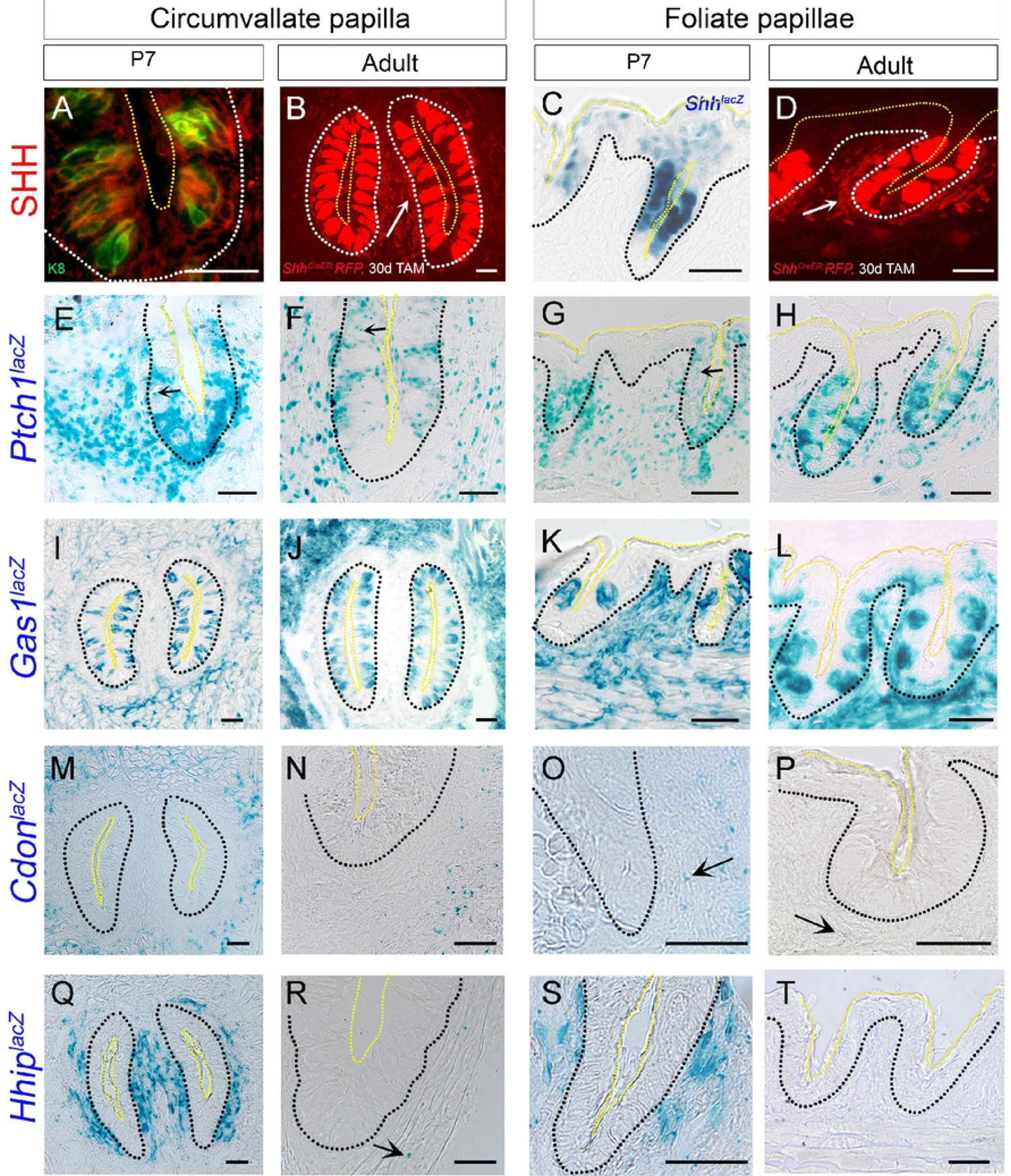
Novel expression of HH pathway receptors in posterior tongue circumvallate and foliate papillae from early postnatal and adult stages. (A-D) SHH ligand expression within CVP and FOP taste bud (TB, labelled with K8, green) as demonstrated by either antibody staining or X-gal staining in *Shh^lacZ^*^/+^ reporter mouse at P7 (A,C) and RFP staining in *Shh^CreER^;RFP* reporter mouse after 30 days of tamoxifen treatment at adult stage (B,D).(E-T) X-gal staining using *Shh^lacZ^*^/+^, *Ptch1^lacZ/+^*, *Gas1^lacZ/+^*, *Cdon^lacZ/+^* and *Hhip^lacZ/+^* reporter mice at P7 and adult stages. In both stages, *Ptch1^lacZ^* is expressed in basal, perigemmal, apical extragemmal and stromal cells of CVP (E,F) and FOP (G,H). Few *Ptch1^lacZ^* puncta is seen within TB of CVP at both stages (E,F, arrows) and FOP at P7 (G, arrow). *Gas1^lacZ^* is present in TB and stromal cells at P7 and in adult CVP (I,J) and FOP (K,L). Stromal expression of *Cdon^lacZ^* is observed in CVP (M,N) and FOP (O,P), but the expression declines at adult stages (N,P) as compared to P7 (M,O). Extensive *Hhip^lacZ^* CVP and FOP stromal expression is observed at P7 (Q,S), which is virtually eliminated by the adult stage (R,T), and few punctate *lacZ*+ cells are observed in CVP stroma at adult stage (R, arrow). Black or white dotted lines outline the epithelium and yellow dotted lines outline the TB apical surface in all the images. Scale bars are 50µm.

**Fig 5.**
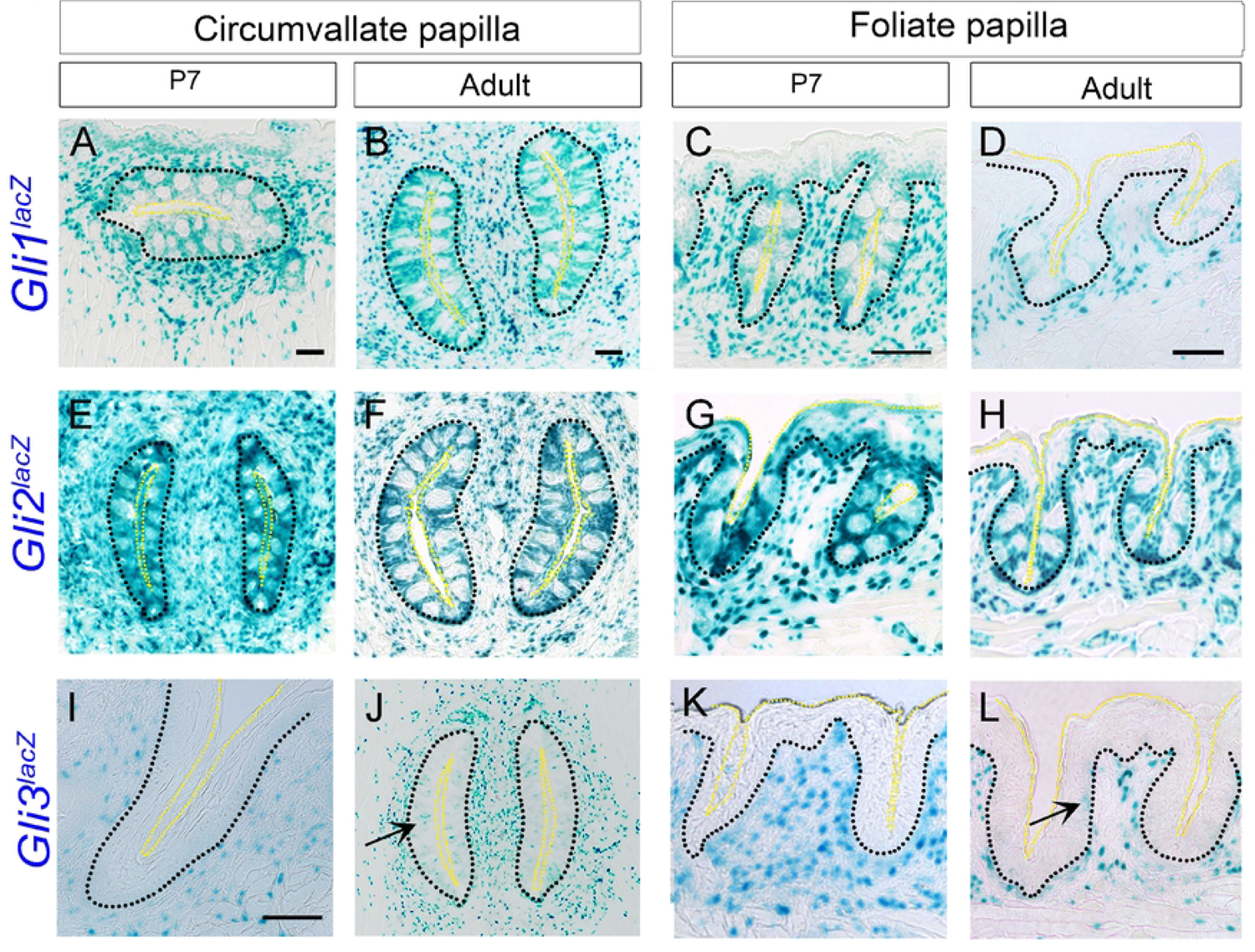
Expression pattern of HH pathway transcription factors in circumvallate and foliate papillae from early postnatal and adult stages. (A-L) X-gal staining of *Gli1^lacZ/+^*, *Gli2^lacZ/+^*and *Gli3^lacZ/+^* reporter mice at P7 and adult stages. *Gli1^lacZ^* (A-D) and *Gli2^lacZ^* (E-H) are expressed in basal, perigemmal, apical extragemmal and stromal cells of CVP (A,B,E,F) and FOP (C,D,G,H) at both the P7 and adult stages. On the other hand, *Gli3^lacZ^* is present in CVP and FOP stromal cells (I-L) at P7 (sagittal section) and adult tongue (horizontal section) and additionally in TB in adult CVP and FOP TB (J,L arrows). Black dotted lines outline the epithelium and yellow dotted lines outline the TB apical surface in all the images. Scale bars (50µm) in A,B,C and D apply to respective column images except I.

For *Ptch1^lacZ^* expression, we investigated CVP and FOP sagittal sections at P7 and horizontal sections at the adult stage (**Fig 4E-H)**. We observed *Ptch1^lacZ^*+ cells in CVP and FOP basal, perigemmal, apical extragemmal and stromal cells at both stages (**Fig 4E-H**). Magnification of the CVP wall shows few punctate *Ptch1^lacZ^* expressions within TB at either P7 or adult stages (**Fig 4E,F**, arrows). Few *Ptch1^lacZ^* are seen within FOP TB at P7 (**Fig 4G**, arrow).

X-gal staining of horizontal sections of the *Gas1^lacZ^* mouse posterior tongue at P7 and adult revealed expression in CVP and FOP TB (**Fig 4I-L**). There is extensive *Gas1^lacZ^* expression in CVP stroma, but stromal cells of CVP adjacent to papilla walls had few *Gas1^lacZ^*+ cells as compared to overall *Gas1^lacZ^* CVP stromal expression at both the stages (**Fig 4I,J**). *Gas1^lacZ^* is expressed throughout FOP stroma (**Fig 4K,L**).

We investigated horizontal CVP and FOP sections of *Cdon^lacZ^* mice at the P7 and adult stages (**Fig 4M-P**). Contrasting with *Gas1*, *Cdon^lacZ^* expression is not seen in either CVP or FOP TB. The *Cdon^lacZ^*+ stromal cell expression pattern at P7 in the CVP connective tissue core is comparable to the *Gas1^lacZ^* arrangement, with few positive cells surrounding papilla walls whereas the expression is prominent in the rest of the stromal cells distant to papilla walls (**Fig 4M**). We observed *Cdon^lacZ^*+ cells throughout FOP stromal cells at P7 (**Fig 4O**). In both CVP and FOP at adult stages (**Fig 4N,P**), the expression declined as compared to P7 (**Fig 4M,O**). Unlike the anterior FP expression pattern (**Fig 1J-L**), we do not observe *Cdon^lacZ^* expression in CVP or FOP basal cells (**Fig 4M-P**).

In contrast to *Gas1^lacZ^* and *Cdon^lacZ^* expression patterns in the connective tissue core*, Hhip^lacZ^* is extensively expressed in CVP and FOP stromal cells neighboring the papilla walls at P7 (**Fig 4Q,S**). Further, expression of *Hhip^lacZ^* is limited to a few stromal cells in adult CVP and virtually absent in the adult FOP connective tissue core, as indicated in horizontal sections (**Fig 4R**, arrow; **4T**).

Overall, the expression patterns of *Shh* ligand and HH membrane receptors *Ptch1, Gas1,* and *Cdon* remain unchanged during the postnatal development of CVP and FOP. Only *Hhip* expression changed in the CVP and FOP connective tissue core at 8 weeks of age as compared to the first postnatal week. Interestingly, epithelial expression of *Gas1* and *Cdon*, as seen in anterior tongue, is not observed in the posterior CVP and FOP at any given developmental stage(s). Further, stromal CVP and FOP *Hhip* expression at an early developmental stage is coincident with expression in FP stroma. The data indicate the requirement of different HH receptors between anterior and posterior tongue epithelial development and further suggest distinct roles of *Hhip* in early and late developmental stages of FP, CVP and FOP stroma.

### Transcription factor *Gli1* and *Gli2* expressions in the posterior circumvallate and foliate papillae remain the same from the first week of birth through eight weeks, while *Gli3* expression is modified

We analyzed horizontal or sagittal CVP and FOP sections of *Gli1^lacZ^* in P7 and adult tongues (**Fig 5A-D**). In both stages, we observed expression in CVP and FOP basal, perigemmal, apical extragemmal and stromal cells (**Fig 5A-D**). Similar to *Gli1^lacZ^, Gli2^lacZ^*+ cells are present in CVP and FOP basal, perigemmal, apical extragemmal and stromal cells throughout postnatal and adult stages (**Fig 5E-H**). There is no *Gli1^lacZ^* or *Gli2^lacZ^* expression within TB of CVP or FOP (**Fig 5A-H**).

In X-gal staining of P7 sagittal tongue sections there are *Gli3^lacZ^*+ cells in CVP and FOP stroma (**Fig 5I,K**). Expression of *Gli3^lacZ^* in CVP and FOP stroma surrounding papilla walls remain similar in adult horizontal tongue sections (**Fig 5J,L**) as compared to P7 (**Fig 5I-K**). In addition, there are positive cells in CVP and FOP TB in mice older than 8 weeks (**Fig 5J,L**, arrows).

The data clearly indicate that expression of HH pathway transcription factors, *Gli1* and *Gli2*, in CVP and FOP mostly remains the same from early postnatal weeks through adult stages, and the location patterns are similar to each other and to the anterior tongue.

### HH signaling component expressions in adult soft palate is similar to fungiform papilla

We investigated the soft palate (SP) with taste organs in the oral cavity that harbor TB distributed in the epithelium, not enclosed in papillae. We limited our investigations to the adult stage. X-gal staining of the *Shh^lacZ^* mouse SP reveals expression in TB (**Fig S1L**). Further, with the 14-day tamoxifen-induced *Shh^CreER^;RFP* mouse model, SHH is observed in both TB and nerves (**Fig 6A**). Different membranous HH receptors have distinct expressions: *Ptch1^lacZ^* in SP perigemmal and stromal cells with some punctate expression within TB cells (**Fig 6B**); *Gas1^lacZ^* in SP basal epithelial, stromal and TB cells (**Fig 6C**); *Cdon^lacZ^* in the SP basal epithelium, mainly in the lower half (**Fig 6D**). We did not observe expression of HH membrane receptor and antagonist *Hhip^lacZ^* in SP (**Fig 6E**) as reported earlier (27). All HH transcription factors *Gli1^lacZ^, Gli2^lacZ^* and *Gli3^lacZ^* are expressed within SP stromal cells (**Fig 6F-H**). *Gli1^lacZ^* and *Gli2^lacZ^* are additionally expressed in SP basal epithelial, perigemmal and apical extragemmal cells (**Fig 6F,G**). Our HH component expression data reveal active HH signaling in SP, similar to FP. We further propose paracrine HH signaling in the SP from TB cells to perigemmal, basal and stromal cells, as reported earlier for the FP (9).

**Fig 6.**
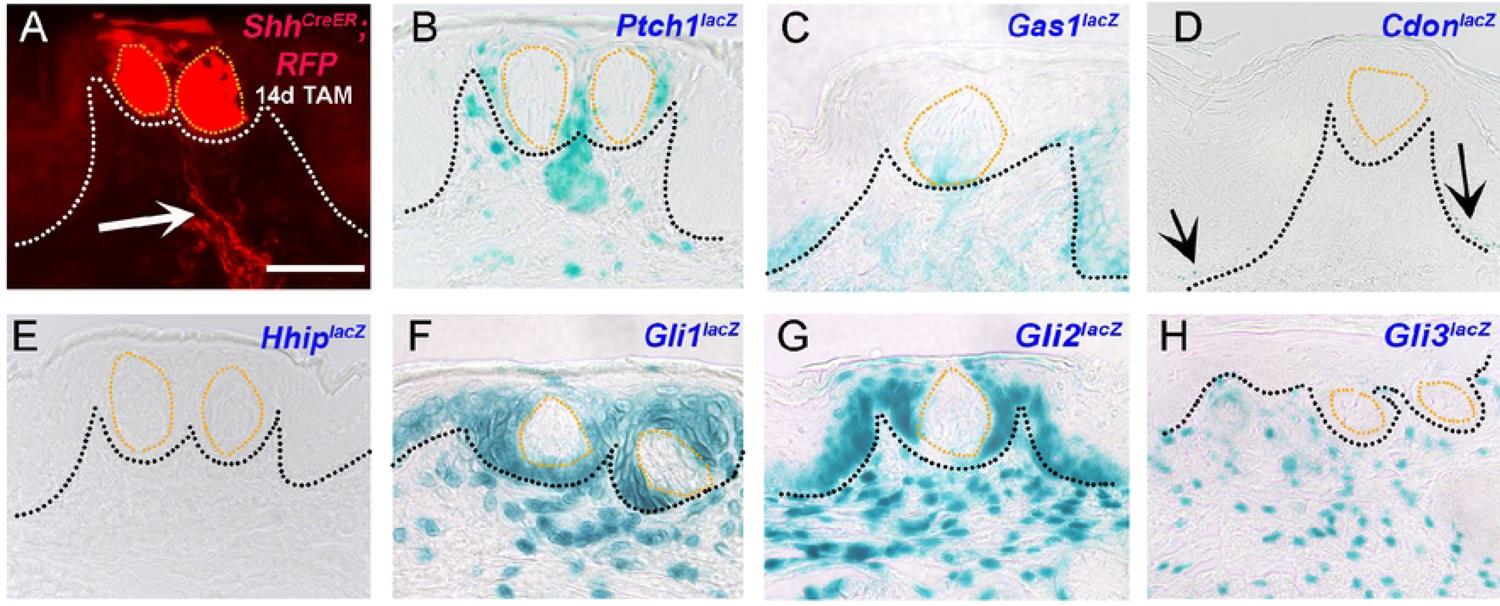
Novel expression of HH pathway components in adult mouse soft palate. (A) RFP staining in *Shh^CreER^;RFP* reporter mouse after 14 days of tamoxifen treatment reveals *Shh* expression in the soft palate (SP) TB and nerves (A, arrow). (B) *Ptch1^lacZ^* is expressed in perigemmal and stromal cells along with punctate expression within TB. (C) *Gas1^lacZ^* is present within the TB, epithelium and stroma. (D) Expression of *Cdon^lacZ^* is seen at the SP epithelium rete ridges (D, arrows). (E) *Hhip^lacZ^* is not expressed in the SP. (F-G) Both *Gli1^lacZ^* (F) and *Gli2^lacZ^* (G) are present in SP basal, perigemmal, apical extragemmal and stromal cells. (H) *Gli3^lacZ^* is exclusively present in the stromal cells. Black or white dotted lines outline the epithelium and orange dotted lines outline the TB in all the images. Scale bar (50µm) in A applies to all images.

### SHH ligand, membrane receptor *Ptch1*, and transcription factors *Gli1* and *Gli2* are the only HH pathway components expressed in adult geniculate and trigeminal ganglia

Geniculate ganglion (GG) and trigeminal ganglion (TG) contain cell bodies of chorda tympani and lingual nerves, respectively, that innervate the anterior tongue (50). We have previously shown the presence of SHH ligand in adult mouse GG and TG using *Shh^CreER^;RFP* mouse model (26). Here, we have used a SHH antibody and reproduced the finding of SHH positive soma in both GG (**Fig 7B,F,J,N**) and TG (**Fig 7D,H,L,P**). Among all the HH co-receptors, *Ptch1* is the only one expressed in both GG and TG (**Fig 7A-D**). Further, most of the *Ptch1^lacZ^*+ cells are associated with SHH+ cells in adult GG and TG (**Fig 7B,D**). There is no expression of *Gas1^lacZ^* (**Fig 7E-H**), *Cdon^lacZ^* (**Fig 7I-L**) or *Hhip^lacZ^* (**Fig 7M-P**) in either GG or TG.

**Fig 7.**
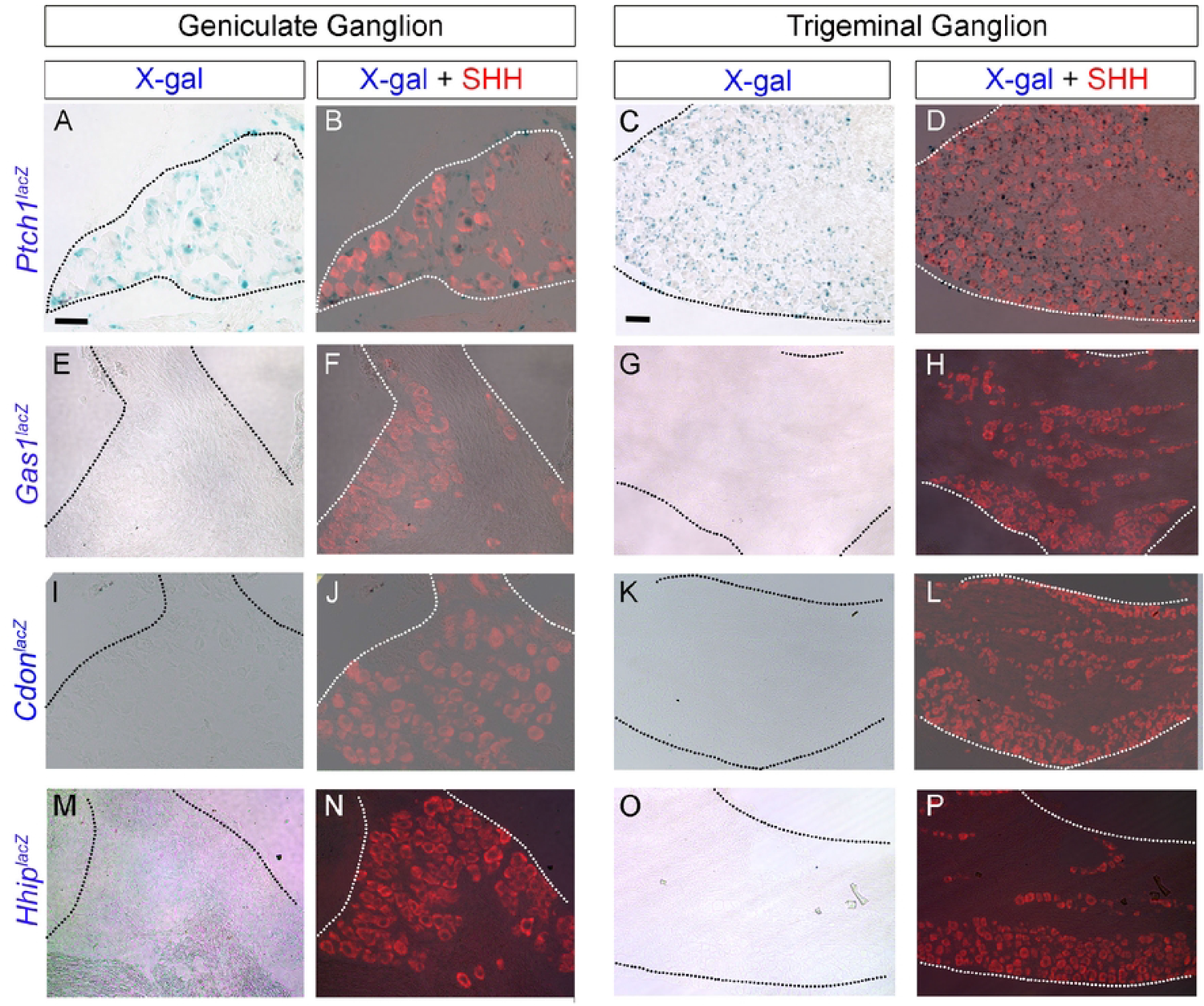
Expression pattern of HH ligand and receptors in adult mouse geniculate and trigeminal ganglia. (A-P) X-gal staining in *Ptch1^lacZ/+^*, *Gas1^lacZ/+^*, *Cdon^lacZ/+^* and *Hhip^lacZ/+^* reporter mice. SHH antibody staining indicates SHH+ (red) cell bodies in GG (B,F,J,N) and TG (D,H,L,P). *Ptch1^lacZ^* expression is observed both in GG (A,B) and TG (C,D) and is predominantly associated with SHH ligand (B,D red). None of the other HH-co-receptors, *Gas1^lacZ^* (E-H), *Cdon^lacZ^* (I-L) and *Hhip^lacZ^* (M-P) are expressed in either GG or TG. Scale bars (50µm) in A and C apply to all Geniculate Ganglion and Trigeminal Ganglion images, respectively.

Between the transcription factors, *Gli1^lacZ^*+ and *Gli2^lacZ^*+ cells but not *Gli3^lacZ^* cells are observed in both GG and TG (**Fig 8A-L**). Further, *Gli1^lacZ^*+ and *Gli2^lacZ^*+ cells are either next to SHH+ cells or seem to be associated with nerve fibers (**Fig 8B,D,F,H**). Importantly, all the SHH+ neurons do not have adjacent *Gli1^lacZ^*+ and *Gli2^lacZ^*+ cells suggesting specificity towards a neuron cell type for a particular function.

**Fig 8.**
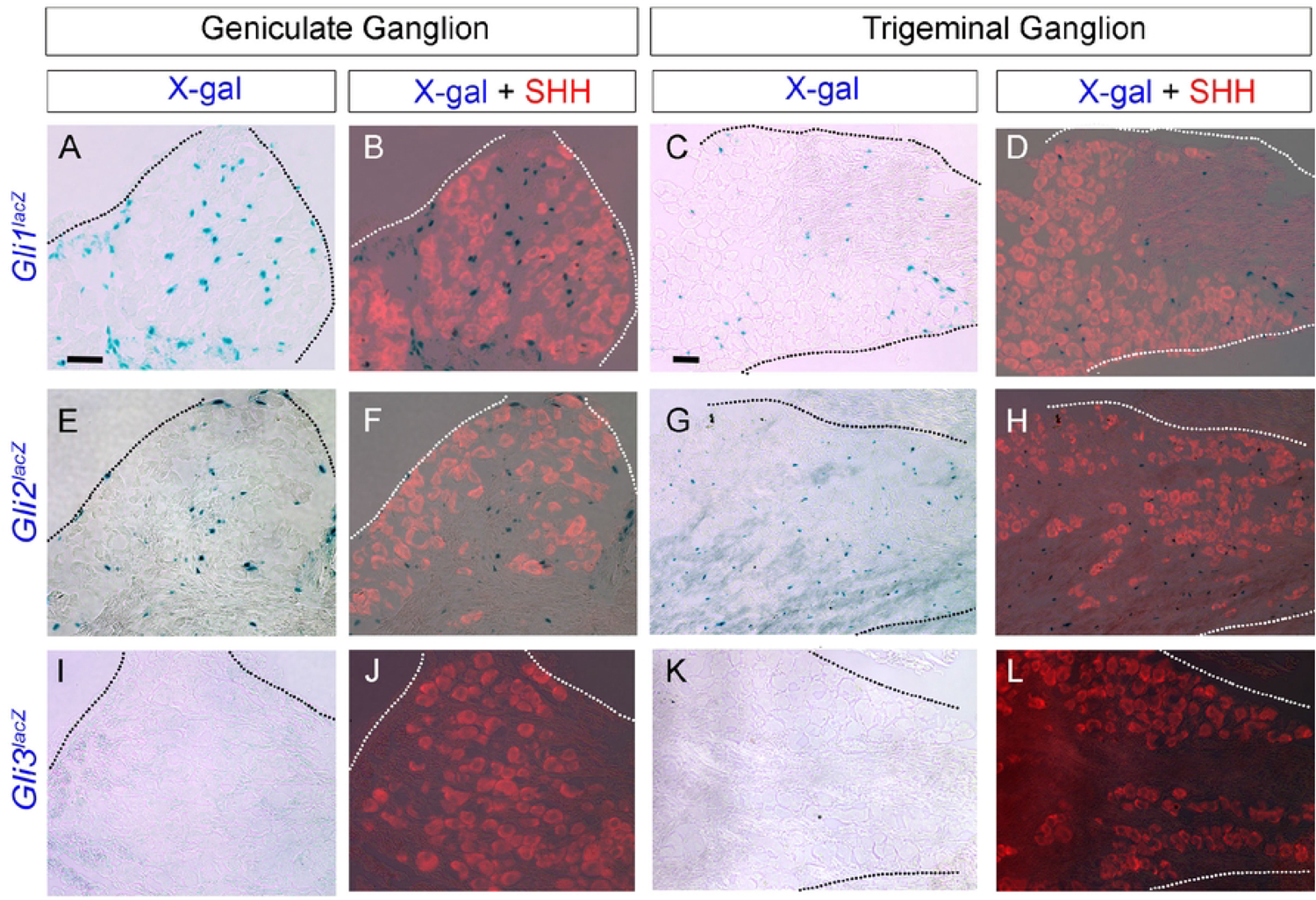
Expression pattern of HH transcription factors in adult mouse geniculate and trigeminal ganglia. (A-L) X-gal staining in *Gli1^lacZ/+^*, *Gli2^lacZ/+^* and *Gli3^lacZ/+^* reporter mice. SHH antibody staining indicates SHH+ (red) cell bodies in GG (B,F,J) and TG (D,H,L). Both *Gli1^lacZ^* (A-D) and *Gli2^lacZ^* (E-H) expression are observed in GG (A,B) and TG (C,D) and predominantly associated with SHH ligand (B,D,F,H red). In contrast, the transcription factor *Gli3^lacZ^* is not expressed in either GG or TG. Scale bars (50µm) in A and C apply to all Geniculate Ganglion and Trigeminal Ganglion images, respectively.

Overall, the presence of SHH, *Ptch1*, *Gli1* and *Gli2* indicates a requirement for active HH signaling in GG and TG while the absence of *Gas1*, *Cdon*, *Hhip* and *Gli3* implies that there is no HH antagonism in both ganglia.

## Discussion

This is the first detailed expression analysis of HH pathway components, namely ligand *Shh*, membrane receptors *Ptch1*, *Gas1*, *Cdon* and antagonist *Hhip*, and GLI transcription factors *Gli1*, *Gli2* and *Gli3,* in the gustatory system during development and/or homeostasis. We have included six distinct tissues, FP, CVP, FOP, SP, GG, and TG, to compare the similarities and differences in their HH signaling component expressions (**Table 1**). Our data unveil intriguing and unique expression patterns of pathway components in the lingual epithelium, stroma or muscles. We established the expression time-course for these components throughout postnatal tongue development and identified shifts in expression during late postnatal stages that match with their conclusion of morphogenesis by postnatal day 21 (10, 15, 16, 18). We provide evidence that the HH pathway receptors, *Gas1* and *Cdon*, that remain unexplored in the tongue, have extensive yet dynamic lingual epithelium, stromal or muscular expressions. Our study opens the opportunity for future functional studies to understand the individual roles of HH pathway components in distinct tissues of the gustatory system.

**Table 1.**
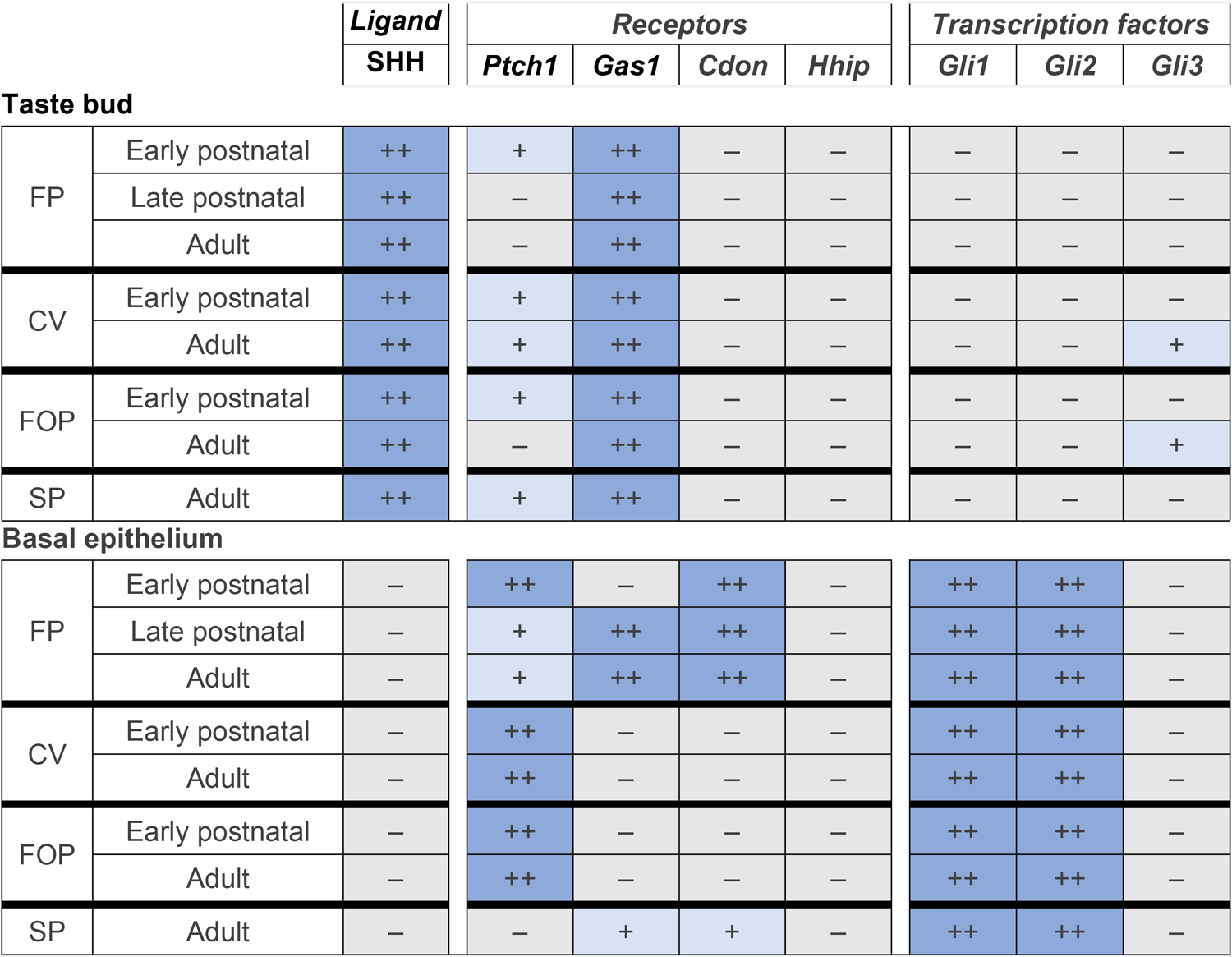
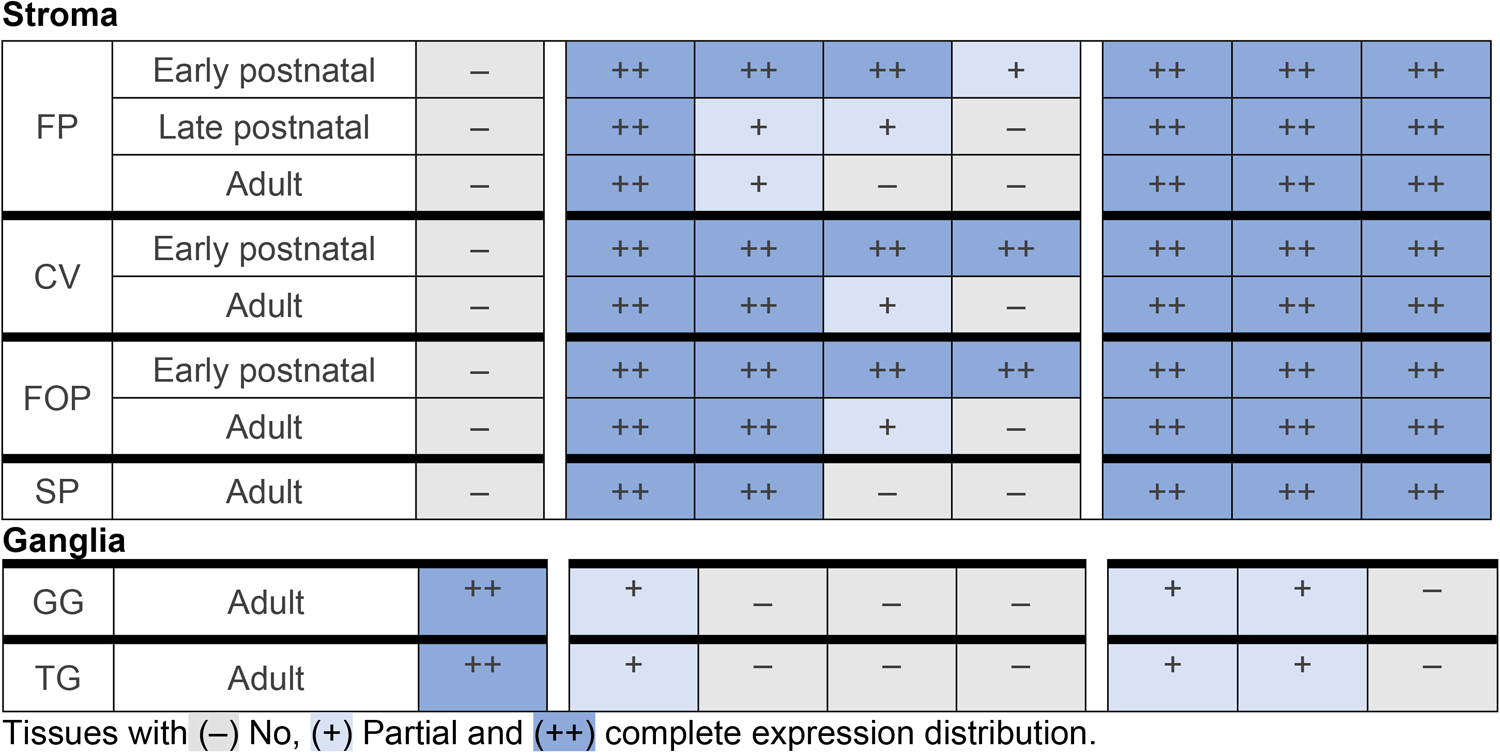
Summary of HH pathway component expressions in gustatory system.

|  |  | *Ligand* | *Receptors* |  |  |  | *Transcription factors* |  |  |
| --- | --- | --- | --- | --- | --- | --- | --- | --- | --- |
|  |  | SHH | *Ptch1* | *Gas1* | *Cdon* | *Hhip* | *Gli1* | *Gli2* | *Gli3* |
| **Taste bud** |  |  |  |  |  |  |  |  |  |
| FP | Early postnatal | ++ | + | ++ | – | – | – | – | – |
|  | Late postnatal | ++ | – | ++ | – | – | – | – | – |
|  | Adult | ++ | – | ++ | – | – | – | – | – |
| CV | Early postnatal | ++ | + | ++ | – | – | – | – | – |
|  | Adult | ++ | + | ++ | – | – | – | – | + |
| FOP | Early postnatal | ++ | + | ++ | – | – | – | – | – |
|  | Adult | ++ | – | ++ | – | – | – | – | + |
| SP | Adult | ++ | + | ++ | – | – | – | – | – |
| **Basal epithelium** |  |  |  |  |  |  |  |  |  |
| FP | Early postnatal | – | ++ | – | ++ | – | ++ | ++ | – |
|  | Late postnatal | – | + | ++ | ++ | – | ++ | ++ | – |
|  | Adult | – | + | ++ | ++ | – | ++ | ++ | – |
| CV | Early postnatal | – | ++ | – | – | – | ++ | ++ | – |
|  | Adult | – | ++ | – | – | – | ++ | ++ | – |
| FOP | Early postnatal | – | ++ | – | – | – | ++ | ++ | – |
|  | Adult | – | ++ | – | – | – | ++ | ++ | – |
| SP | Adult | – | – | + | + | – | ++ | ++ | – |
| Stroma |  |  |  |  |  |  |  |  |  |
| FP | Early postnatal | – | ++ | ++ | ++ | + | ++ | ++ | ++ |
|  | Late postnatal | – | ++ | + | + | – | ++ | ++ | ++ |
|  | Adult | – | ++ | + | – | – | ++ | ++ | ++ |
| CV | Early postnatal | – | ++ | ++ | ++ | ++ | ++ | ++ | ++ |
|  | Adult | – | ++ | ++ | + | – | ++ | ++ | ++ |
| FOP | Early postnatal | – | ++ | ++ | ++ | ++ | ++ | ++ | ++ |
|  | Adult | – | ++ | ++ | + | – | ++ | ++ | ++ |
| SP | Adult | – | ++ | ++ | – | – | ++ | ++ | ++ |
| Ganglia |  |  |  |  |  |  |  |  |  |
| GG | Adult | ++ | + | – | – | – | + | + | – |
| TG | Adult | ++ | + | – | – | – | + | + | – |

Tissues with (–) No, (+) Partial and (++) complete expression distribution.

### Ligand SHH is present in taste bud and nerves of tongue and soft palate

While previous studies have demonstrated that SHH is expressed in FP taste bud cells and nerves (9, 26, 45, 47, 51), data on other taste organs are lacking and focused mainly on the adult stage. Using X-gal reporter mouse models, here we show that *Shh* ligand is consistently expressed within the taste buds of FP, CVP and FOP at both postnatal and adult stages (**Table 1**). In addition, using a tamoxifen-inducible transgenic reporter for *Shh*, we additionally observe *Shh* in the nerves entering TB of CVP, FOP and SP. Nerve-derived HH ligands participate in touch dome (52, 53) and hair follicle (54) maintenance. In contrast, when taste nerve-derived *Shh* was deleted in mouse, there were no effects on postnatal or adult FP taste cell differentiation (8, 45). In our nerve-cut experiments, both taste and lingual, in adult mouse, we observed a reduction in the number of taste buds (27), which could be due to the additional loss of other taste nerve-derived growth factors. The effects of neural SHH deletion on posterior tongue and SP taste buds have not yet been evaluated. Recently, it has been shown that misexpression of *Shh* in lingual epithelium induces ectopic TB formation in FILIF and the number of ectopic TB is 3-fold higher when overexpression occurs at P14 as compared to P1 (8). It is not clear what changes are occurring in HH signaling regulation during early postnatal stages that lead to different outcomes of *Shh* alteration.

### HH signaling is broad in anterior lingual epithelium during postnatal tongue development

HH signaling in adult tongue is restricted to taste organs (27). However, HH-responding *Gli1*+ cells are expressed in both taste FP and non-taste FILIF in the initial postnatal weeks (**Fig 3A,B**). For postnatal stages, a previous study utilized *in situ* staining and showed *Gli1* expression in FP only (9). However, *in situ* staining might have limitations for detecting lower gene expression (55). Thus, we utilized *Gli1* reporter mouse models and confirmed *Gli1*+ cells in FP and intriguingly revealed additional expression in FILIF. Importantly, *Gli1*+ cells at two different locations of a tissue performed distinct differentiation and proliferation functions in the mandible (56) and cranial region (57). Whether *Gli1*+ cells in FP and FILIF have different sets of HH target genes to perform distinct functions remains to be investigated.

FILIF grow rapidly in the first postnatal week and steadily thereafter until P21, when they reach full maturation (58). HH/SMO/GLI inhibition at adult stage altered FP but did not alter FILIF structure and pattern (27). However, the function of HH signaling in FILIF development is yet to be defined. Our data suggest that HH signaling regulates FILIF maturation and after the conclusion of postnatal morphogenesis by P21, FILIF becomes HH-independent.

### Novel membrane HH co-receptors are identified in tongue taste and non-taste papillae epithelium

We recently reported the core HH membrane receptor *Ptch1* expression in FP and anterior epithelial face of FILIF and suggested dual roles for the receptor: activator in FP and antagonist in FILIF (27). In addition to PTCH1, there are additional co-receptors, GAS1, BOC and CDON, that help create the SHH gradient during organ development (34, 35, 59–63) but remain unknown in the tongue. Overlapping roles for *Gas1* and *Cdon* were demonstrated by upregulating the Shh target transcriptional response (19). Here, we have unmasked the overlapping expression patterns of *Gas1* and *Cdon* in the late postnatal lingual epithelium, including both FP and FILIF.

Initial studies claimed *Gas1* as an antagonist of HH signaling (59, 64) while subsequent studies showed positive regulation by *Gas1* (19, 20, 63). Remarkably, *Gas1* is not expressed in the lingual epithelium before P21 and thus does not overlap with HH signaling activity. Further, HH signaling gets restricted to FP with concurrent expression of *Gas1* in the postnatal lingual epithelium. Thus, we propose that *Gas1* might has dual roles as positive regulators of the signaling in FP and negative regulators in FILIF where there is no SHH ligand.

*Cdon* expression remains the same in the anterior lingual epithelium throughout, irrespective of the changes in *Gli1* and *Gas1* expression. Thus, *Cdon* may act as a multi-functional regulator of HH signaling during tongue maturation by first promoting HH signaling in FILIF and later inhibiting it together with *Gas1*. It might also be possible that *Gas1* and *Cdon* have independent roles in tongue development. Cooperation between *Gas1* and *Cdon* was not observed during limb development (19), even though they are expressed in overlapping domains in the limb (62, 63). Recently, it has been shown that embryonic *Cdon* deletion does not affect HH signaling; however, when combined with *Gas1* deletion, it leads to the elimination of HH signaling (60). Co-expression of *Gas1* and *Cdon* in FILIF along with simultaneous reduction of HH signaling suggests the latter antagonistic roles may be dependent on the concurrent presence of *Gas1*. Our data further indicate that *Cdon* does not overlap with *Ptch1* suggesting that SHH employs *Cdon, Gas1* and *Ptch1* in a context-dependent manner during early and late postnatal stages of tongue development. Intriguingly, *Gas1* and *Cdon* are not expressed in the CVP and FOP epithelium, suggesting their different roles in anterior and posterior tongue epithelium.

Further, unlike other receptors, there is novel and extensive expression of *Gas1* in the TB of all three taste papillae. Whether *Gas1* helps in SHH dispersion and functions similarly in FP, CVP and FOP TB remains to be investigated. However, our findings point towards autocrine HH signaling in TB, in addition to paracrine signaling earlier suggested in FP (9). As *Gas1* growth arrest function can be ligand independent (65), we propose that *Gas1* may have distinct functions in TB, FP and FILIFP cell contexts depending on ligand availability.

### Novel membrane HH co-receptors are differentially expressed in tongue stroma

Previously, expressions of *Ptch1*, GLI transcription factors, *Gli1*, *Gli2* and *Gli3* were reported in lingual stromal cells in adults (9, 25, 26, 48, 49). Here, we report an intriguing extensive expression of *Gas1* and *Cdon* in the anterior tongue stroma at P7 that decreases after tongue maturation by P28 and remains low thereafter (**Figs 2G-L, S1C-G**). *Gas1* and *Cdon* are also expressed in posterior CVP and FOP stroma at early postnatal tongue development stage. While *Gas1* expression in CVP and FOP stroma was maintained at adult stage, *Cdon* stromal expression was downregulated in FOP (**Table 1**) suggesting differential roles for *Gas1* and *Cdon* in FP, CVP and FOP stroma.

*Cdon* expression in embryonic lingual mesenchyme was first noted 20 years before (66) but remains uninvestigated. In limb bud, *Cdon*+ HH responding cells extend long specialized filopodia and may participate in relaying activation of the pathway at a distance from the cell soma (21). Remarkably, we observed vimentin+ cells with filopodia extensions to contact the basal lamina (25), but whether they are *Cdon*+ is not known. HH signaling in the embryonic lingual stroma is involved in the transmission of information from epithelial cells to myogenic progenitor cells to coordinate tongue formation (67, 68). Whether these interactions are maintained during postnatal tongue maturation remains to be investigated. We hypothesize that stromal HH signaling via *Gas1* and *Cdon* might regulate lingual epithelium and muscle development in the early postnatal stages.

Intriguingly, we observed antagonist and HH co-receptor *Hhip* expression in the posterior tongue stroma at P7, which was virtually eliminated in the adult tongue. Further, the opposing expression patterns of *Hhip* with *Gas1* and *Cdon* (**Fig 4I,M,Q**) suggest a discrete population of stromal cells, neighboring and distant to the CVP and FOP walls.

### *Gas1* is the only HH pathway component identified in lingual muscles at early postnatal stage

HH signaling is essential for embryonic myoblast migration (69, 70). Here we show that neither SHH nor HH signaling is present in the postnatal muscles. In that case, direct control of HH seems less likely or can be mediated by HH-responsive stroma (67, 68). Importantly, we unmask *Gas1* expression in postnatal lingual muscles at early postnatal stages (**Figs 2G, S1C**) when lingual muscles are in the maturation phase (71, 72). Our recent studies revealed that *Gas1* deletions did not alter embryonic myoblast migration but caused defects in muscle differentiation leading to altered muscle arrangement (73). However, their role in the postnatal tongue is not yet known.

In Down’s syndrome patients (74) and mouse models (75), hyperplastic, disorganized and atrophic neuromuscular junctions in the tongue were seen. Recently, it has been shown that chromosome 21 (which causes Down’s syndrome) transcribes genes that can modulate SHH signaling (76). Although the genes that modulate SHH signaling are yet to be investigated, there is a possibility that perturbed HH signaling can lead to lingual muscle disruption in early postnatal stages. We propose that *Gas1* can have cell-autonomous effects on lingual muscles in early postnatal stages and once muscles are mature, they become HH independent at late postnatal stage.

### Distinct expression of HH signaling components in anterior and posterior tongue at adult stage

As reported in previous studies, we observed *Shh* within TB including basal cells in the three distinct taste papillae. Importantly, though, our data here reveal many important points regarding HH signaling activity in the papillae of adult anterior and posterior tongue (**Fig 9**). We demonstrated the novel presence of the HH-receptor, *Gas1*, within TB in FP, CVP and FOP suggesting that *Gas1* might play a role in SHH dispersion from basal cells to throughout the TB for taste cell differentiation. In addition to *Gas1*, we observed *Gli3* within CVP and FOP TB as reported by another study as a negative regulator of a taste cell type (49), but not in FP (**Fig 9**). Recently, we identified and discussed differences in regulation of the anterior versus posterior tongue (50).

**Fig 9.**
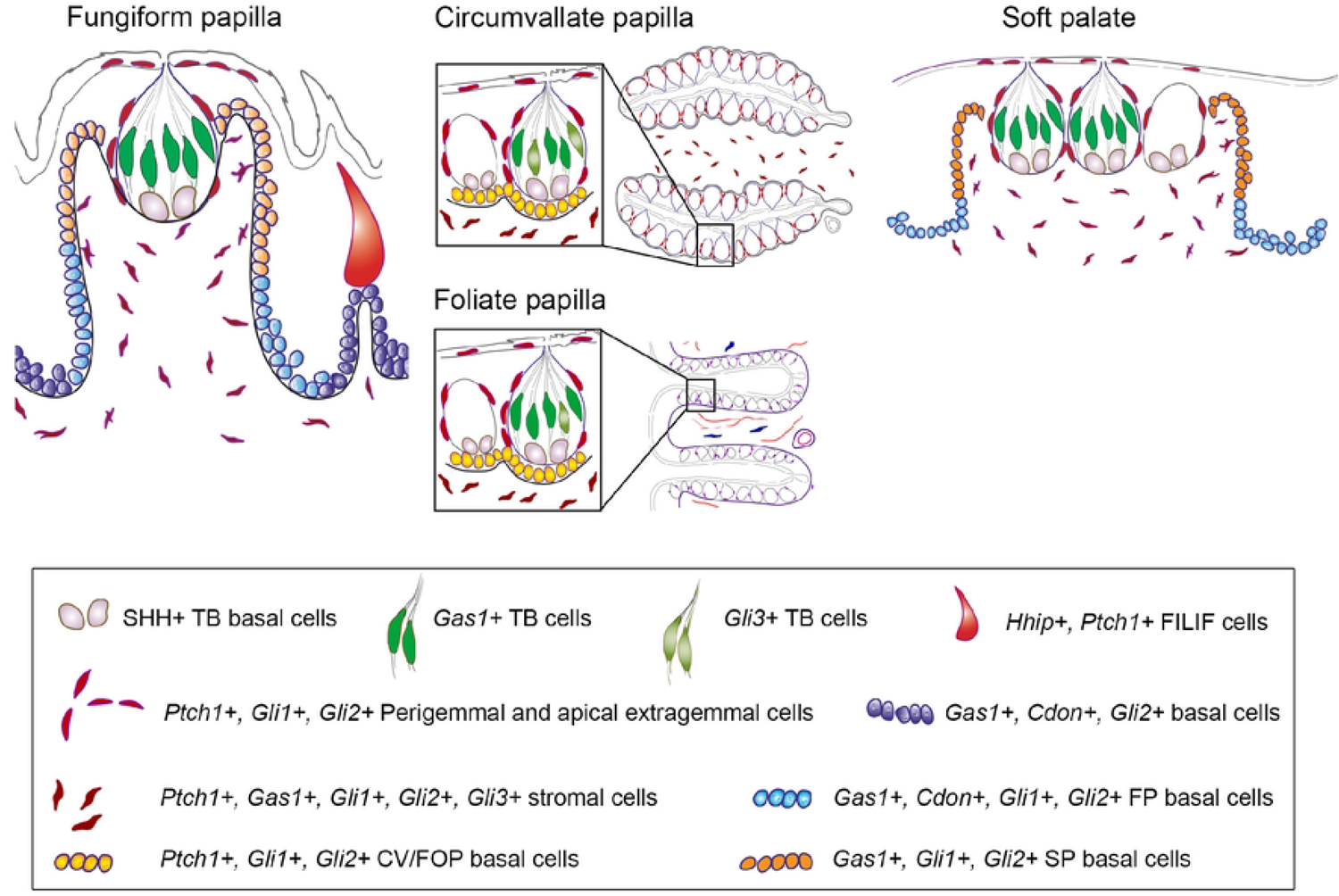
Expression pattern of HH signaling components in the adult FP, CVP, FOP and soft palate. Building from on Fig 1A, HH signaling components are located in FP, CVP, FOP and soft palate. SHH is predominant in TB basal cells. *Gas1* is present in all papillae TB, whereas *Gli3* is also expressed in the posterior tongue papillae TB. *Hhip* and *Ptch1* are present in anterior epithelial face of FILIF. *Gli1*+, *Ptch1*+ and *Gli2*+ cells are observed in lingual basal, perigemmal, apical extragemmal and stromal cells of all the papillae. However, *Ptch1* is primarily present in the upper apical half of FP. On the other hand, HH co-receptors *Gas1* and *Cdon* are concentrated in lower half of FP wall and entire lingual epithelium. *Gas1* and *Gli3* are also expressed in the stromal cells of all the papillae and soft palate. The expression pattern of HH signaling components in the soft palate is similar to that observed in FP.

Current findings further emphasize the anterior-posterior tongue differences, as indicated by *Gas1* and *Cdon* expression in the anterior lingual basal epithelium but not in the posterior epithelial cells (**Fig 9**). The data indicate requirement of *Gas1* and *Cdon* to create a gradient to restrict HH signaling to adult FP in anterior tongue. As entire CVP and FOP basal epithelium is HH responsive, formation of HH gradient is not required and thus *Gas1* and *Cdon* do not play any role in posterior tongue epithelium.

Even though *Ptch1*, *Gli1* and *Gli2* are present in both anterior and posterior lingual basal epithelium, *Ptch1* expression pattern varies in the anterior FP as compared to CVP and FOP in adult (**Fig 9**). While the *Ptch1*+ cells from FP basal cells get restricted to the apical half of FP, the entire CVP and FOP basal cells continue to express it. Similar to FP, a decline in *Ptch1* expression is also observed in the dental inner enamel epithelium to maintain the SHH-responsive quiescent adult stem cells (77). Our previous studies emphasized alterations in apical basal proliferation (25, 26, 28) and elimination of *Ptch1* (27) after HH pathway inhibition, implying that *Ptch1* controls FP apical cell proliferation. While the basal half of FP retained cell proliferation, it could not prevent TB loss (25, 26, 28), which further suggests FP stem cell maintenance by *Ptch1*+ cells. The data indicate heterogeneous cell populations in the FP basal epithelium, which demarcates boundaries between the apical and basal epithelium (**Fig 9**).

Stromal expression of all the components is observed but whether they are co-expressing remains to be investigated. We propose stromal diversity in the context of HH signaling components to create distinct niche described previously (78). HH antagonist *Hhip* is specific to non-taste FILIF and is not observed in any of the taste papillae epithelium (**Fig 9**).

### HH signaling in other oral tissue at adult stage

In addition to FP, TB are also housed in the soft palate epithelium without being enclosed in a papilla (**Fig 9**, soft palate). The maturation of soft palate TB is almost complete by 1 week of age and is much faster than that of the TB of FP (10). Upon analyzing soft palate TB in adult mice, we found expression patterns similar to FP (**Fig 9**). Co-expression of *Shh* and *Gas1* in the TB suggests autocrine signaling. The expression patterns of the HH receptors *Ptch1* and *Gas1* and all three *Gli* transcription factors in mesenchyme corroborate a previous study conducted between embryonic days 13 and 14.5 (79). In contrast to their embryonic soft palate data, we observe epithelial expression of *Ptch1*, *Gas1*, *Gli1* and *Gli2* and no expression of *Hhip* in the adult soft palate. Whether there is a change in expression of HH components as noted in our FP expression studies or a limitation of the ^35^S-in situ hybridization technique (79) remains to be investigated. *Cdon* was not studied by them, and we observe an expression pattern in the epithelium similar to *Gas1*. It might be possible that *Gas1* and *Cdon* are positive regulators of HH signaling in the soft palate.

We show that HH-responding *Gli1*-cells are present in the soft palate epithelium, and that epithelium gives rise to TB (80). When we treated adult rats with the HH pathway inhibition drug sonidegib, we observed that TB were reduced to half their normal numbers and the remaining TB were atypical (28) suggesting that HH signaling regulates adult soft palate TB. Embryonic studies showed roles for soft palate epithelial Shh in mediating mesenchymal interactions controlling palatal outgrowth (67, 81–83). Our studies suggest paracrine signaling regulation from *Shh*+ TB and nerves to HH-responding palatal epithelium and stroma.

### HH signaling in ganglia associated with gustatory system during adulthood

Multimodal chorda tympani and somatosensory lingual nerves receive afferents from the geniculate (GG) and trigeminal ganglion (TG), respectively (84). SHH expression in the GG and TG marks the ligand sources of the HH signal and was present in all the neurons (26, 45). As a result, we see *Shh*+ nerves in FP and soft palate that receive chorda tympani nerve fibers. The expression of *Shh* in lingual nerve is not clear even though TG express SHH+ neurons (28, 46, 47).

Here, we reveal the expression patterns of other members of HH signaling (**Table 1**). Among the HH receptors, only *Ptch1* is expressed in both GG and TG in association with SHH+ cells. The HH antagonist *Hhip* is not expressed in either of the ganglia. Transcription factors *Gli1* and *Gli2* are expressed in GG and TG, predominantly next to SHH+ cells. Importantly, there are many SHH+ neurons that do not have associated expression of the transcription factors (**Fig 8**). Our data suggest that HH signaling regulates GG and TG and requires components different than the tongue or the soft palate. The functions of HH-responsive GG and TG remain to be investigated.

Our comparative analysis of unexplored HH pathway co-receptors in anterior and posterior tongue, soft palate and ganglia, in epithelium, stroma or muscles offers insight into HH pathway activity in distinct organs of the gustatory system. Future functional studies can reveal the regulatory functions of heterogeneous expression patterns within distinct tongue compartments. Our studies lay the foundation for future functional studies to further dissect the role of HH signaling components in postnatal tongue tissues development.

## Acknowledgements

We would like to acknowledge Dr. Chen-Ming Fan for sharing the *Gas1^lacZ^* mouse model with AK.

## Supporting information

**Supplementary Fig 1. Expression pattern of HH signaling components in the anterior and posterior tongue and soft palate.** (A,B) X-gal staining in *Ptch1^lacZ/+^* reporter mouse indicates a reduction in *Ptch1*+ FP basal cells and TB cells at P19 (A) and expression in FILIF anterior epithelial face at P7 (B). (C-E) X-gal staining in *Gas1^lacZ/+^* reporter mouse at P4, P21 and adult stages suggests that while lingual muscles express *Gas1^lacZ^* at P4, *Gas1*+ muscle cells are not observed at P21 and adult. Ecad antibody co-staining reveals absence of *Gas1^lacZ^* expression in lingual epithelium at P4 (C). Epithelial *Gas1^lacZ^* expression is observed at P21 (D) and maintained through the adult stage (E). (F,G) X-gal staining in *Cdon^lacZ/+^* reporter mouse reveals stromal expression at P8 (F), which gets downregulated at adult stage (G). K8 antibody co-staining confirms no *Cdon ^lacZ^* expression in TB (F). (H,I) X-gal staining in *Hhip^lacZ/+^* reporter mouse shows stromal *Hhip^lacZ^* expression at P12 in tongue (H) and below the TB (H, inset, arrow) but not at P30 (I). (J) X-gal staining in P14 *Gli1^lacZ/+^* reporter mouse indicates reduced *lacZ* expression in FILIF (arrow). (K,L) X-gal staining in *Shh^lacZ^* reporter mouse shows expression in the taste buds of CVP at P7 (K) and soft palate at adult stage (L). Black dotted lines outline the epithelium. Scale bars are 50µm.

## Notes

### Competing Interest Statement

The authors have declared no competing interest.

